# Optimization of negative stage bias potential for faster imaging in large-scale electron microscopy

**DOI:** 10.1101/2020.09.03.277830

**Authors:** Ryan Lane, Yoram Vos, Anouk H. G. Wolters, Luc van Kessel, Ben N.G. Giepmans, Jacob P. Hoogenboom

**Author notes:** Correspondence, Ryan Lane, Jacob Hoogenboom.

## Abstract

Large-scale electron microscopy (EM) allows analysis of both tissues and macromolecules in a semi-automated manner, but acquisition rate forms a bottleneck. We reasoned that a negative bias potential may be used to enhance signal collection, allowing shorter dwell times and thus increasing imaging speed. Negative bias potential has previously been used to tune penetration depth in block-face imaging. However, optimization of negative bias potential for application in thin section imaging will be needed prior to routine use and application in large-scale EM. Here, we present negative bias potential optimized through a combination of simulations and empirical measurements. We find that the use of a negative bias potential generally results in improvement of image quality and signal-to-noise ratio (SNR). The extent of these improvements depends on the presence and strength of a magnetic immersion field. Maintaining other imaging conditions and aiming for the same image quality and SNR, the use of a negative stage bias can allow for a 20-fold decrease in dwell time, thus reducing the time for a week long acquisition to less than 8 hours. We further show that negative bias potential can be applied in an integrated correlative light electron microscopy (CLEM) application, allowing fast acquisition of a high precision overlaid LM-EM dataset. Application of negative stage bias potential will thus help to solve the current bottleneck of image acquisition of large fields of view at high resolution in large-scale microscopy.

## Introduction

Mapping the full ultrastructural layout of complex biological systems at nanometer-scale resolution is a major challenge in cell biology. Electron microscopy (EM) is uniquely capable of stretching the vast spatial scales necessary to identify macromolecular complexes, subcellular structures, and intercellular architecture. As a consequence, interest in large-scale EM, where many high-resolution tiles are stitched into a gigapixel image frame, has exploded in recent years. Large-scale EM, however, suffers from the long acquisition times necessary to acquire sufficient signal at high resolution (Peddie and Collinson, 2014).

A variety of approaches have been undertaken to advance throughput. While throughput is already a bottleneck for large-scale 2D imaging (Kuipers et al., 2015), most of these approaches have been developed under the framework of 3D imaging. Throughput is particularly relevant to the field of connectomics in which it typically takes months to acquire the image data necessary for neuronal reconstruction (Kornfeld and Denk, 2018). To image the brain of a larval zebrafish, for example, Hildebrand et al. (2017) conducted multiple imaging rounds at successively higher magnification. Regions of interest (ROI) were selected between imaging rounds for successive, targeted acquisitions down to 4 nm/px resolution, thereby reducing the time it would otherwise take to fully image the full brain at high resolution. Similarly, Delpiano et al. (2018) used detection of in-resin preserved fluorescence in an integrated light and electron microscope for automated guiding to ROIs for subsequent acquisition. Other approaches involve parallelizing the imaging load across multiple instruments. This has been employed in focused ion beam scanning electron microscopy (FIB-SEM) for the reconstruction of thick slices of *Drosophila* brain tissue at isotropic (8 nm × 8 nm × 8 nm) voxel resolution (Hayworth et al., 2015) as well as in serial section transmission electron microscopy (ssTEM) for the yearlong acquisition of a cubic millimetre of mouse brain tissue (Yin et al., 2019). Dedicated instrumentation for faster imaging of serial thin sections has also been developed in recent years. In some instances conventional microscopes have been equipped with specialized detection optics to allow for larger fields of view (Zheng et al., 2018). Multi-beam instruments in which a sample is simultaneously imaged by multiple focused electron beams have also been developed (Eberle et al., 2015; Ren and Kruit, 2016).

Faster imaging could also be achieved by increasing signal collection in established thin sections approaches, which would allow for reduced acquisition time while maintaining a sufficient signal to noise ratio (SNR). It has previously been shown that the use of a retarding field increases SNR in SEM (Phifer et al., 2009), but for biological imaging the use of a retarding field has thus far been investigated in detail only for serial blockface scanning EM (SBF-SEM) (Bouwer et al., 2016; Ohta et al., 2012). Relatedly, the use of a positive stage bias has been examined for the suppression of secondary electrons (SEs) (Xu et al., 2017). The full benefits of stage bias remain underutilized because optimization criteria and signal detection in a magnetic immersion field, in particular, have yet to be addressed.

In the cases in which a negative bias potential has been used, a voltage is applied to the stage while the pole piece of the electron microscope is kept at ground such that an electric field is generated between the specimen and detector planes. While the primary electron beam experiences a deceleration, the signal electrons experience an acceleration from the specimen towards the dedicated detector (Fig. 1). The ensuing acceleration results in an increase to the collected signal (Šakić et al., 2011) and—if the detector geometry, landing energy, and potential bias are tuned properly— can be used to filter out secondary electrons (Bouwer et al., 2016). The same signal can in principle then be obtained with a shorter acquisition time.

**Fig. 1.**
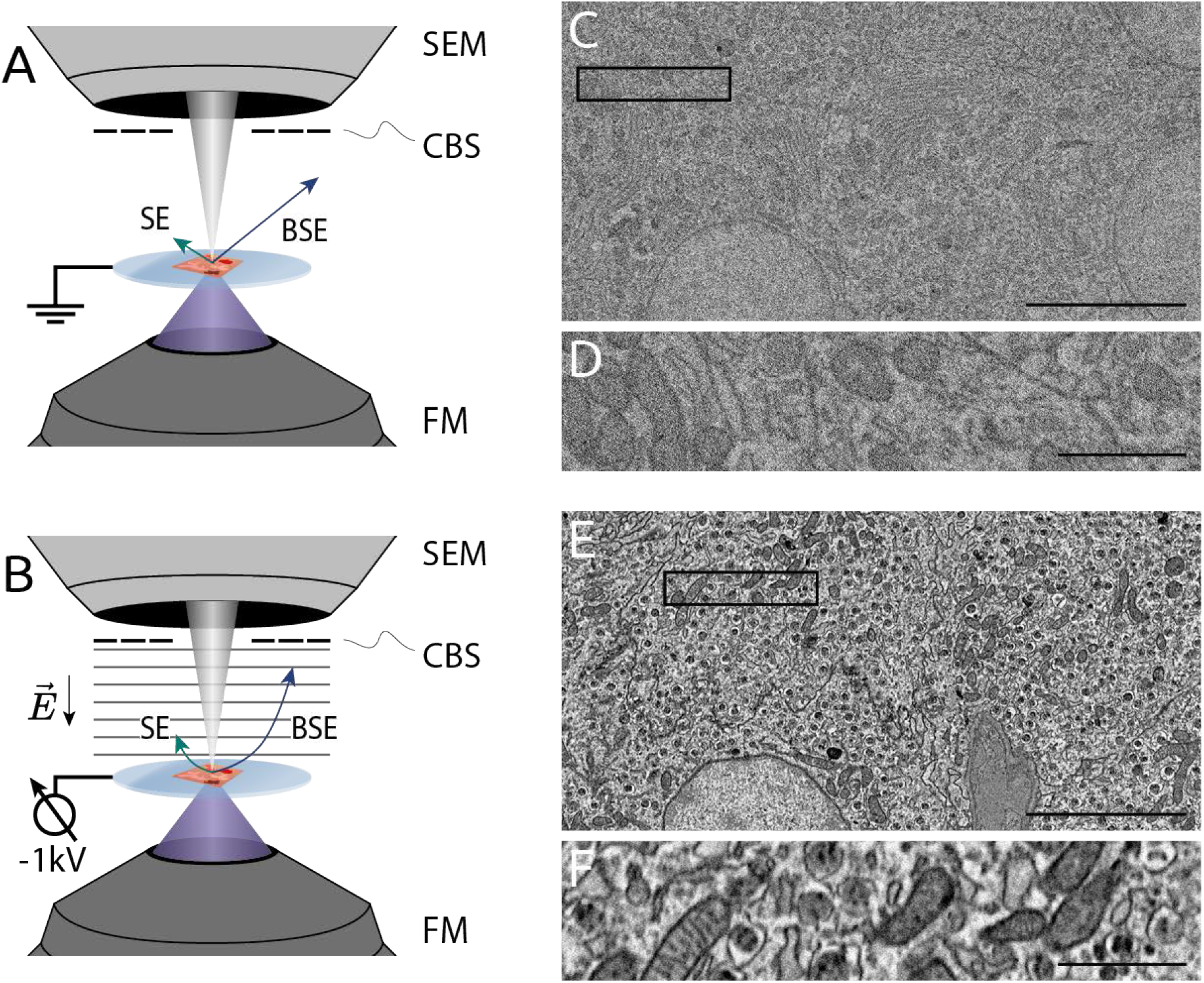
Negative bias potential significantly enhances EM contrast in tissue. Schematic of integrated microscope without (A) and with (B) an applied stage bias. Electric field induced by the bias potential accelerates electrons emitted from the sample to the CBS detector. EM images of rat pancreas tissue without the use of stage bias (C & D). By comparison, EM images of the same tissue acquired with a −1 kV bias potential (E & F). When activating the stage bias, the primary beam energy is increased to maintain a fixed landing energy of 1.5 keV. The per-pixel dwell is held constant at 5 μs. Vast improvement in EM signal and contrast can be seen by comparing insets (D) and (F). Scale bars: 5 μm (C & E); 1 μm (D & F). Raw data at full resolution is available at www.nanotomy.org.

Identification of biological structures and molecules in large-scale EM is typically complemented with approaches to label and visualize specific biomolecules or organelles. Aside from immuno-EM and genetically-encoded enzymatic tags that can deposit osmiophilic polymers, CLEM is perfectly suited to identify entities across spatial scales (reviewed in (de Boer et al., 2015)). However, for serial-section CLEM, preserving fluorescence within the tissue sections requires protocols in which the samples are prepared with limited concentrations of heavy metal staining—and in particular no post-staining to prevent quenching of fluorophores (Kuipers et al., 2015). The reduced amount of staining material then needs to be countered by increased dwell time, further necessitating optimization of EM signal collection.

## Results

### Negative bias potential enhances signal in routine EM samples

We first illustrate how the use of a potential bias can improve signal collection in a typical SEM experiment. A potential bias of −1 kV is applied to the stage of an SEM with an integrated fluorescence microscope (Fig. 1A & B). The bias is applied via an external power supply connected to a custom stage plate such that the sample is electrically isolated from the rest of the fluorescence microscope and electrical components of the stage (Vos et al., 2020). By generating an electric field between the sample and the BSE detector, the bias potential accelerates signal electrons inwards away from their otherwise linear trajectories. Because of their lower energy, secondary electrons (< 50 eV) are redirected inside the inner annulus of the BSE detector, while higher energy backscatter electrons (> 50 eV) are redirected over a wider area depending on their initial emission angle and energy.

Pancreas tissue was prepared for integrated fluorescence-electron microscopy as described in section 6.2. Of note is that for the sake of fluorescence preservation, no post-staining was applied resulting in lower contrast relative to other EM sample preparation protocols (Kuipers et al., 2015). EM images of epon-embedded, 80 nm tissue were acquired in immersion mode with and without a −1 kV bias potential (Fig. 1A & B). When subject to a bias potential, EM images demonstrate higher contrast and less noise (Fig. 1D & F). The primary beam energy was increased by 1 kV such that the landing energy was held constant at 1.5 keV in accordance with the section thickness. Maintaining a fixed landing energy at this magnitude is crucial as it is responsible for the penetration depth of the primary electrons. The 0.4 nA beam current and 5 μs per-pixel dwell time were also held constant between acquisitions. The gain of the BSE detector had to be decreased while applying the negative bias to prevent the detector from saturating.

### Simulating signal electron trajectories with and without negative potential bias

Electron trajectories were simulated to better ascertain how a negative bias potential may give rise to better signal detection. Secondary electron (SE) and backscatter electron (BSE) trajectories were simulated for a variety of EM imaging conditions (Fig. 2). A model of the optical layout within the integrated microscope was developed in Electron Optical Design (EOD) (Lencová and Zlámal, 2008) incorporating the geometry of the FEI ThermoFisher Verios SEM objective lens and concentric backscatter (CBS) detector. The negative potential bias is factored into the model by implementing the sample plane as an additional lens element, which can then be biased to an arbitrary voltage. To mirror the 5 mm working distance of our microscope, the end of the pole piece (grey element in Fig. 2) and start of the sample plane (red) are situated at *z* = 0 mm and *z* = 5 mm respectively. The roughly 0.5 mm thick CBS detector (blue) is then located immediately below the pole piece.

**Fig. 2.**
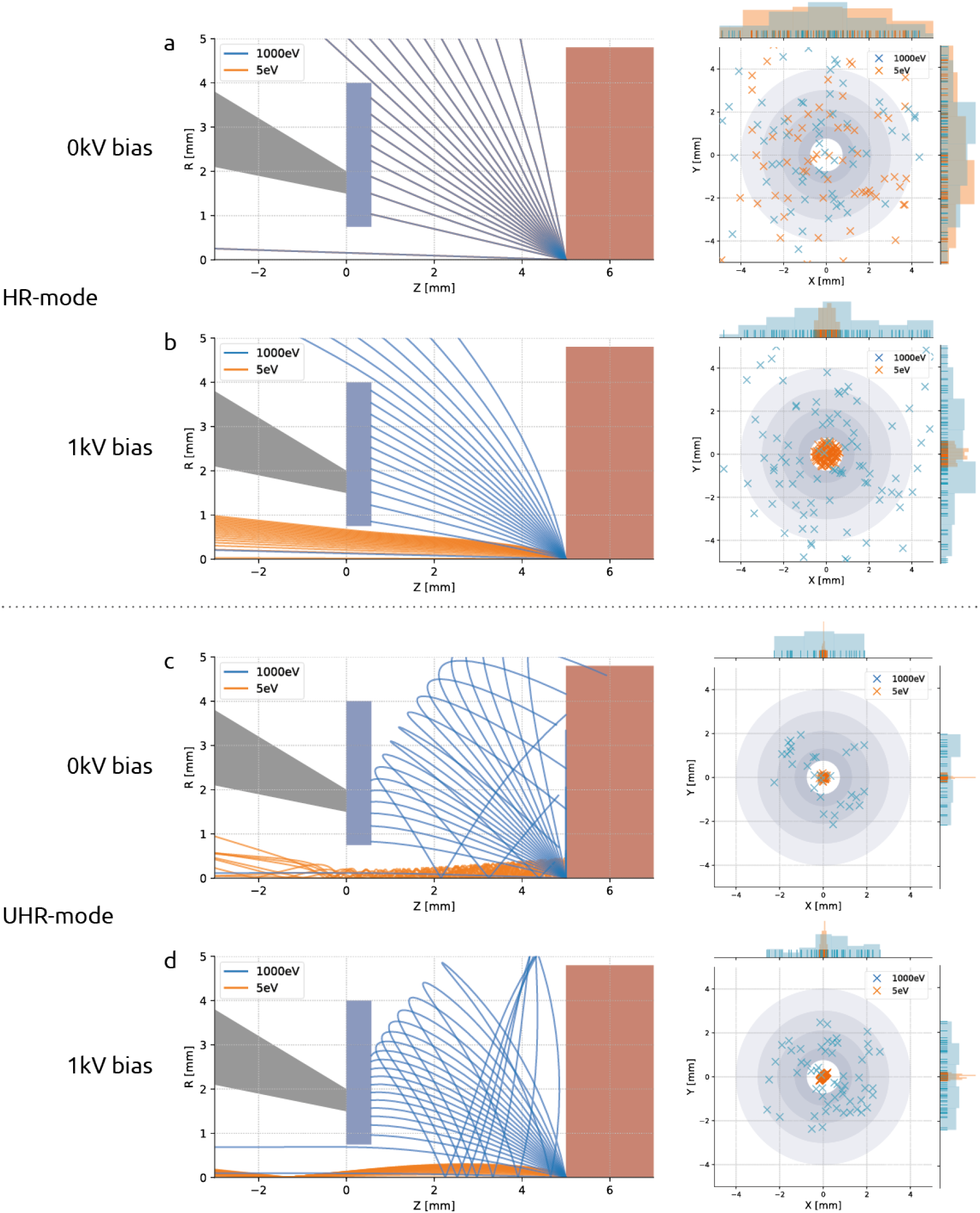
Signal electron trajectories demonstrate the efficacy of stage bias in redirecting BSEs to the detector while simultaneously filtering out secondary electrons. Trajectory plots for SE and BSE bundles launched from the sample plane (left) and scatter plots (right) show the spatial distribution of signal electrons at the detector plane. In HR-mode, SEs and BSEs travel in overlapping, linear paths without the presence of an electric field (a), but BSEs get accelerated towards the detector when a negative bias potential is introduced (b). Signal electrons take on spiral trajectories in the presence of an immersion magnetic field (c), but are again steered to the detector when an electric field is added (d). In each set of simulations, BSEs (blue) and SEs (orange) are launched from the sample plane at *z* = 5 mm. Trajectory plots show geometry of the pole piece (grey), CBS detector (blue), and stage plate (red). Scatter plots show *x, y* coordinates of signal electrons at the detector plane (*z* = 5 mm). Spatial distributions of signal electrons are plotted on the margins of the scatter plots.

Simulations were done for both non-immersion (high resolution or HR) and immersion (ultra-high resolution or UHR) SEM operation modes. For the case of non-immersion mode (Fig. 2A & B), the magnetic focusing field is contained within the objective lens and therefore does not play a role in the signal electron trajectories. In these instances, the trajectories of the SEs and BSEs are dictated entirely by their initial velocity and the electric field due to the bias potential. In UHR-mode (Fig. 2C & D), however, the sample is immersed in a strong magnetic field that both focuses the primary beam and— together with the electric field—alters the paths taken by the signal electrons. For this reason, the magnetic field strength is calculated by the field strength required to focus a parallel beam propagating in the +z direction at the sample plane.

For each scenario shown (Fig. 2), a bundle of secondary (*E*_0_ = 5 eV) and backscatter electrons (*E*_0_ = 1 keV) is emitted from the origin at *z* = 5 mm. The angular distribution is given by Lambert’s cosine distribution (section 6.1; (Reimer, 1998)). A screen is placed at the detector plane to record the radial position of the signal electrons, from which the scatter plots are generated (Fig. 2). The grey rings of varying diameter represent the individual segments of the CBS detector. For the case of non-immersion mode and no potential bias (Fig. 2A), the region between the detector and sample planes is field-free and the signal electrons travel freely in straight paths coinciding with one another. Only when a bias potential is added (Fig. 2B) do the higher energy backscatter electrons diverge from the secondaries, which, due to their low initial energy, are accelerated inside the BSE detector before they are able to spread out radially. The trajectories change when under the influence of a magnetic immersion field (Fig. 2C) in which case the Lorentz force causes the signal electrons to spiral about the optical axis (MÜLLEROVÁ and KONVALINA, 2009). The low energy SEs remain tightly coiled as they propagate up through the BSE detector while the higher energy BSEs stretch out over greater radial distances. Whether the BSEs collide into the detector depends largely on the emission angle. The addition of a 1kV bias potential (Fig. 2D) enables BSEs with a wider distribution of emission angles to reach the detector, resulting in the collection of more signal. These results suggest no secondary electron is ever registered as a count by the BSE detector—either because it is accelerated inside the detector or (in the field-free case) because it is of too little energy to generate an electron-hole pair (Šakić et al., 2011).

The collection of BSEs increases monotonically with increasing negative bias potential for both imaging modes (Fig. 3). These results agree with what is suggested by the trajectory plots of Fig. 2—that the electric field generated by the stage bias tapers the radial spread of the backscatters leading to a greater percentage of BSEs collected. Note that the percentage of BSEs detected is greater for HR-mode across the range of bias potentials simulated. It therefore seems advantageous to prefer non-immersion mode, however, greater collection efficiency is only one factor to consider. The magnetic immersion field results in lower aberrations, meaning that for high resolution imaging, UHR-mode is still often favourable.

**Fig. 3.**
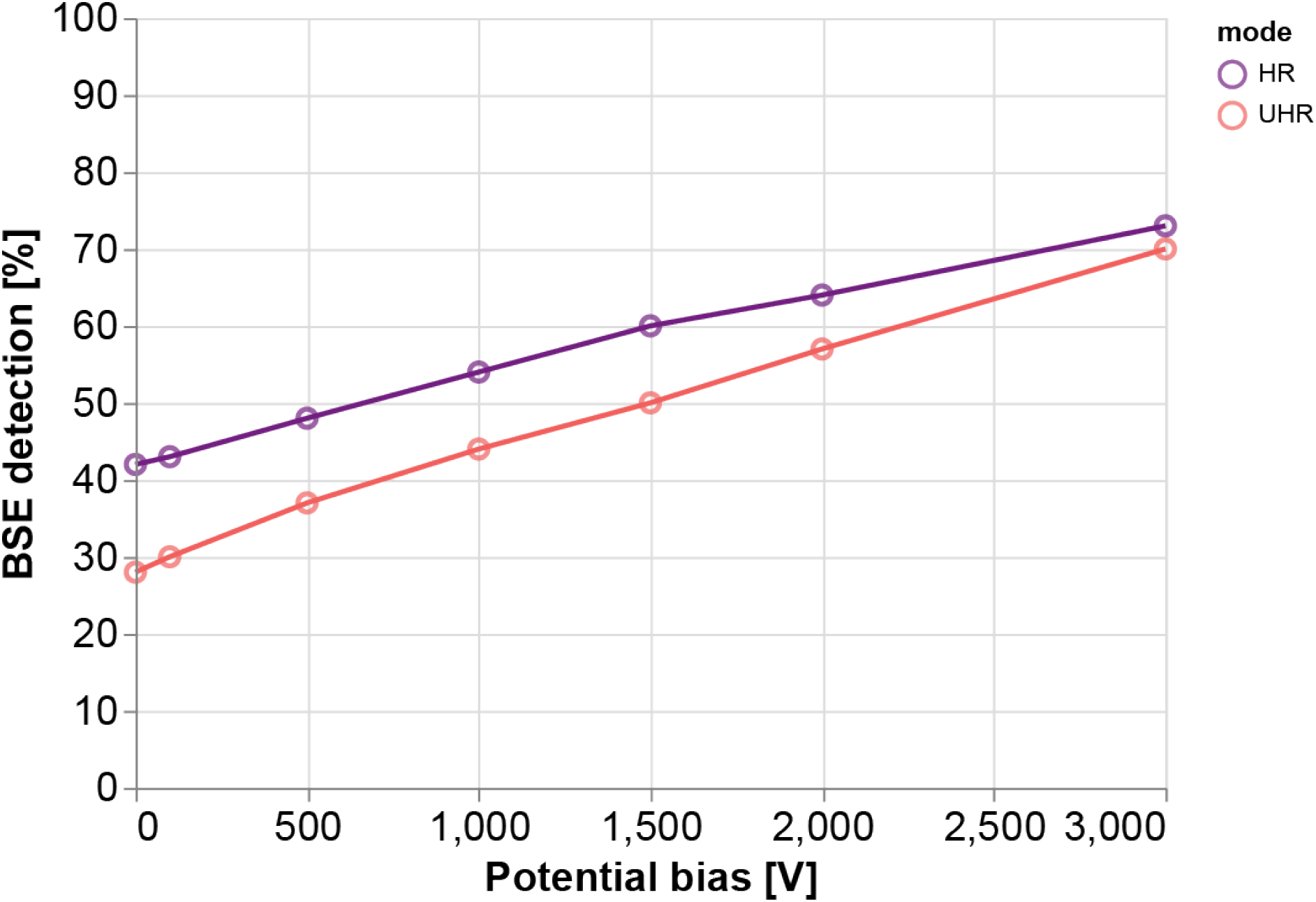
Detection of backscattered electrons is projected to double over the full range of simulated bias potentials. BSE detection rates are derived from simulation trajectories in which the number of electrons incident on the detector is aggregated for each imaging mode and bias voltage. The collection efficiency nearly doubles as the bias is increased from 0 V to −3 kV in non-immersion mode and more than doubles in immersion mode. Simulated bias voltages did not exceed −3 kV as this was the maximum voltage allowed by our experimental setup due to high voltage considerations. The upward trend is, however, not infinite as BSEs with smaller emission angles would eventually be accelerated up through the inner annulus of the detector—along with the SEs—for sufficiently high bias voltages.

### Experimental optimization of negative potential bias leads to increased throughput

EM imaging was expanded to encompass a wider imaging parameter space across a sequence of dwell times and negative bias potentials for both immersion and non-immersion mode based on the simulations (Fig. 4). The primary beam energy was increased together with the bias potential to hold the landing energy constant at 1.5 keV. Likewise, the gain of the CBS detector was adjusted with each bias potential to keep the intensity levels from clipping. The detector gain and offset were manually calibrated to acquire over the full 16-bit range of the detector. This was not always possible, however, as many of the images acquired with low or no bias potential took up only a fraction of detector’s bandwidth—even at maximum gain.

**Fig. 4.**
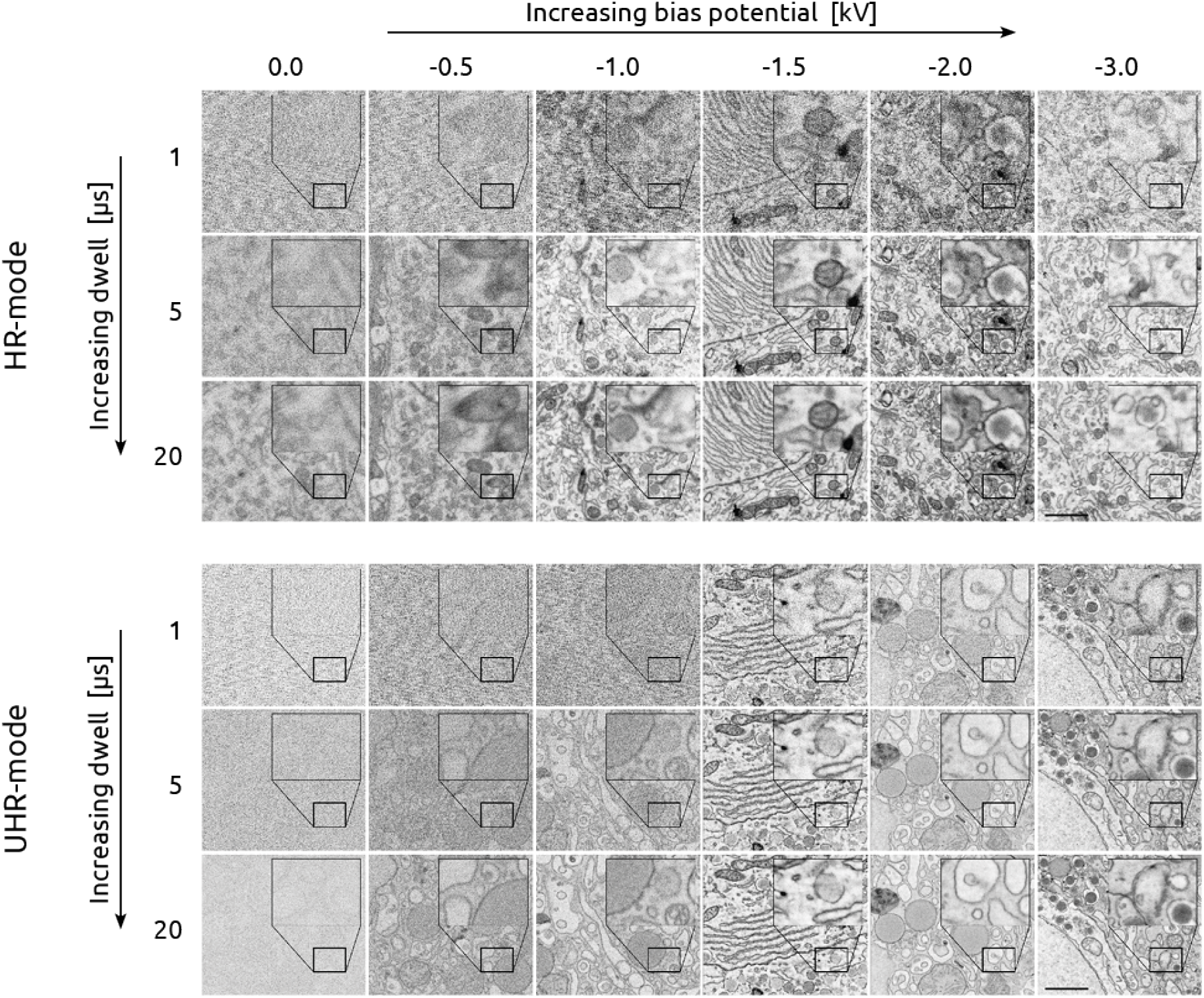
Negative bias potential delivers 20 times faster imaging while maintaining image quality. Bias potential varies from 0 to −3 kV (left to right) while the integration time varies from 1 – 20 μs (top to bottom) for both the non-immersion (top) and immersion mode (bottom) image matrices. All images acquired with 1.5 keV landing energy to match penetration depth. Scale bars: 1μm. Raw data is available at www.nanotomy.org.

An increase in image signal with increasing negative bias potential for both imaging modes up to roughly −1500 V was recorded (Fig. 4), after which it becomes difficult to perceive notable differences in image quality. The signal appears to improve more gradually in non-immersion mode, whereas the improvement for immersion mode is more abrupt. Furthermore, in many instances, increasing the integration time by more than an order of magnitude results in a less substantial increase to the apparent SNR than a 500 V increase to the negative bias potential. This is significant as the integration time is typically the primary imaging parameter to improve image quality—and large increases come at the direct expense of throughput.

Quantitative signal-to-noise measurements were made (Fig. 5) on the collection of images and averaged for each combination of bias potential, dwell, and imaging mode, using the calculation method presented in (Joy, 2002) and corroborated using a method based on the image power spectral density (PSD) (Sup. Fig. 7). In particular, these measurements reveal that an image acquired in non-immersion mode with a 1 μs per-pixel dwell time and −1.5 kV bias potential yields roughly the same SNR as an image acquired with a 5 μs dwell but with no applied bias. The effect of the potential bias is even more pronounced in immersion mode where the SNR of a 1 μs px^-1^ image with a 1.5 kV stage bias exceeds that of a 20 μs px^-1^ image acquired without a bias.

**Fig. 5.**
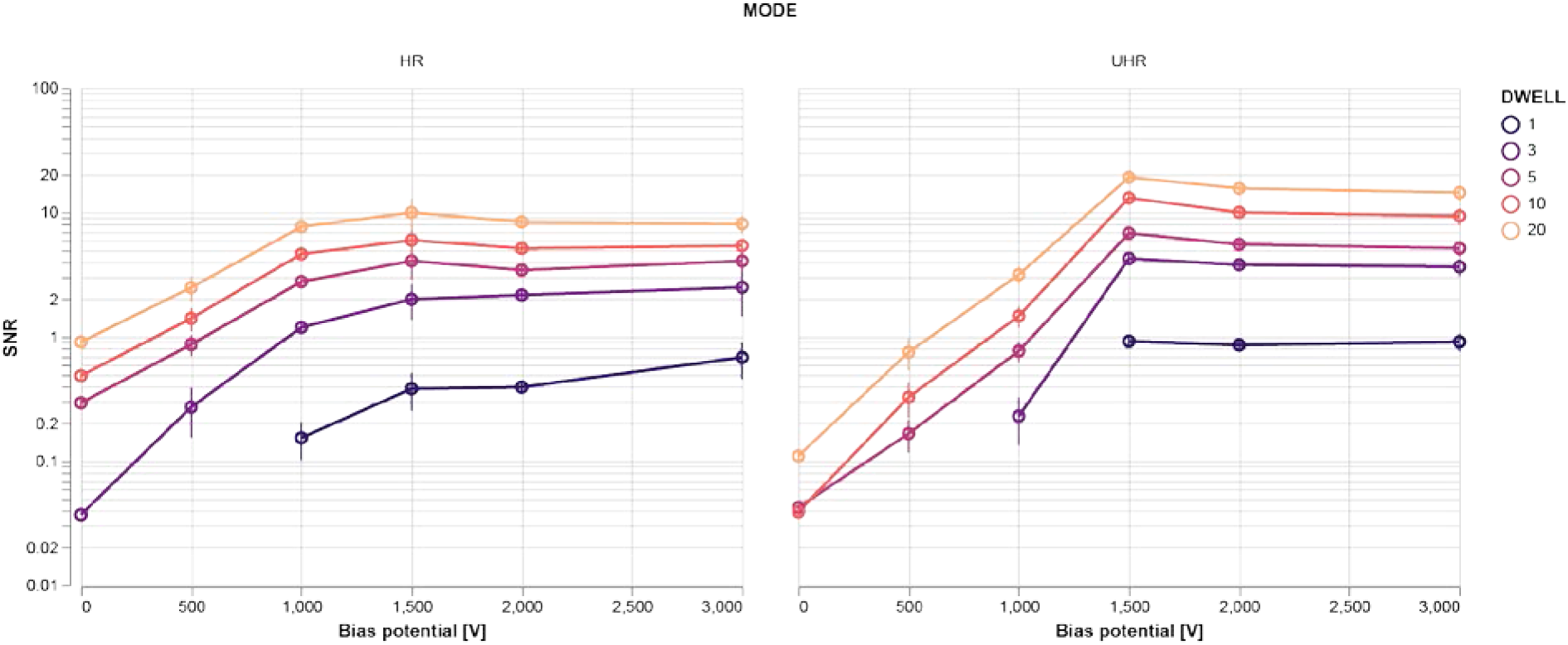
Optimization of bias potential delivers SNR increases of multiple orders of magnitude. At bias potentials greater than 1.5 kV, the SNR is found to level off for both imaging modes. Images are comprised of varying stage bias potentials, integration times, and imaging modes but with fixed 1.5 keV landing energy and 5 mm working distance. Different color lines represent different dwell times as indicated by the legend. SNR measurements are averaged over five EM images at different areas of the tissue for each combination of bias potential, dwell time, and imaging mode. Error bars indicate the standard deviation in the SNR over the five images.

### Potential bias allows for higher throughput EM and CLEM acquisitions

Only small regions of interest are typically recorded at high resolution EM given that full section imaging at sub-10 nm resolution often takes an excessive amount of time. As a result of the enhanced signal-to-noise ratio afforded to us by the use of a negative bias potential, we are able to significantly expedite the imaging of a full tissue section at 5 nm resolution (Fig. 6). Full section (~0.5 mm^2^) acquisition including fluorescence imaging, stage translations, and additional overhead factors was completed in 8 hours. Based on our empirical results (2.3), a negative potential bias of −1.5 kV was chosen for EM imaging in immersion mode. A per-pixel dwell time of 2 μs was chosen to balance high SNR and image clarity with overall imaging time. We note that no post-staining was applied to this section in order to allow integrated acquisition of fluorescence for high-precision overlaid FM.

**Fig. 6.**
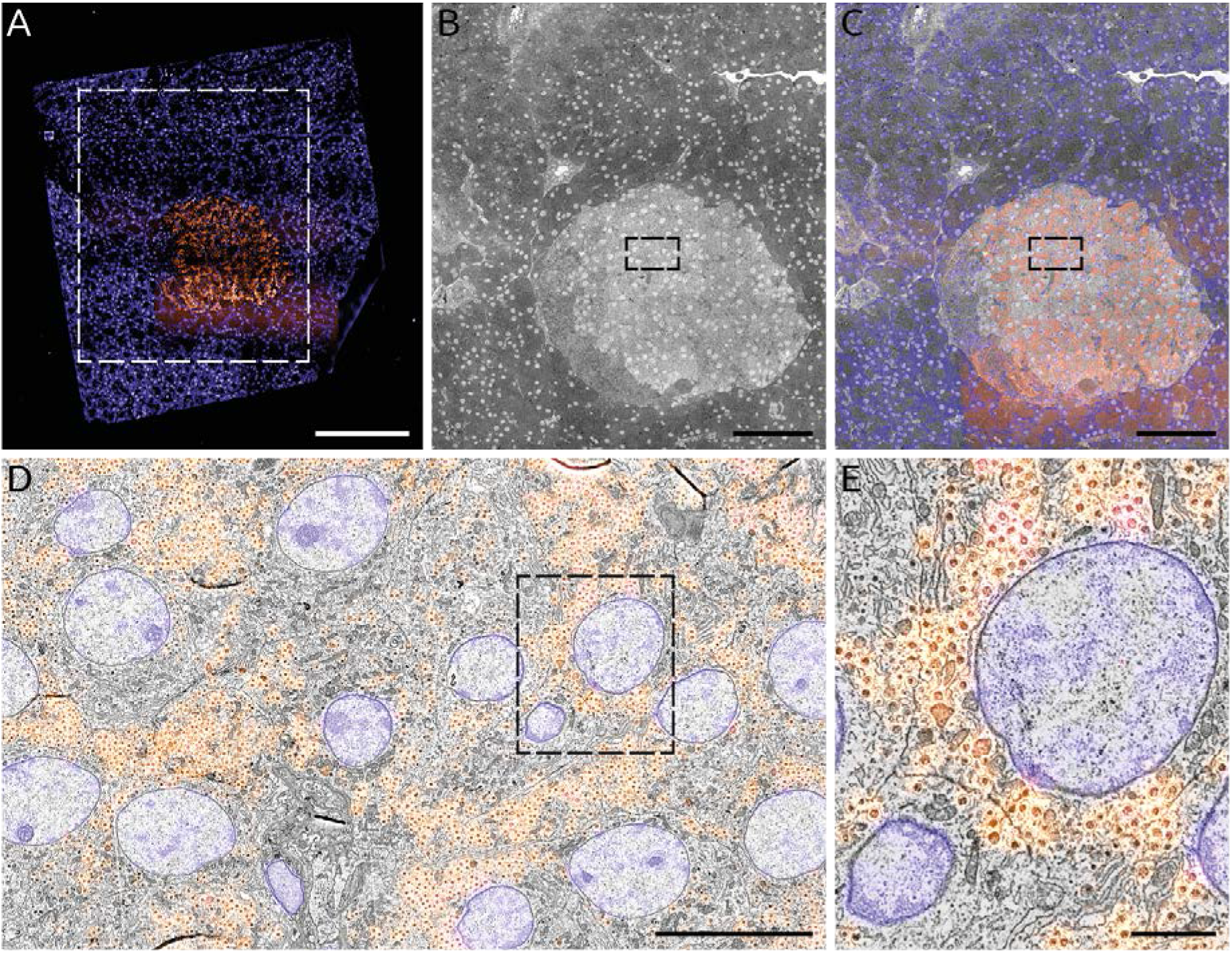
Fast, correlative imaging of a complete EM section at high resolution. 80 nm rat pancreas tissue was imaged at 3 keV beam energy with a −1.5kV stage bias (1.5keV landing energy) with 2 μs dwell as a nanotomy map. (A) Composite two-channel FM image of the tissue section: cell nuclei (blue) stained by Hoechst; insulin-producing beta cells (orange) immunolabeled with Alexa 594. (B) Composite EM image of the area outlined in (A) comprising the islet of Langerhans identified via FM imaging. (C) Correlative overlay of the islet and surrounding exocrine tissue. (D) Zoomed-in area of islet outlined in (B & C) with inset (E) exhibiting the native resolution (5 nm pixel size) that exists across the entirety of the nanotomy map. Total imaging time is 8 hr, the majority of which is taken up by the high-resolution EM imaging. Note that a similar area at this pixel size (see e.g. (Ravelli et al., 2013)) typically takes upwards of 24 hrs with TEM. Scale bars: 200 μm (A); 100 μm (B & C); 10 μm (D); 2 μm (E). Raw data is available through www.nanotomy.org.

Fluorescence images were acquired prior to EM to prevent quenching of the fluorescence due to electron beam irradiation. The insulin-producing beta cells—clustered within the islet of Langerhans— were immunolabelled and given a Hoechst counterstain to target cell nuclei as well as the rough endoplasmic reticulum in the exocrine region of the tissue (blue) (Fig. 6). The section edges can easily be discerned from the FM images, facilitating the area selection for subsequent EM imaging (Fig. 6B). Here the islet (light grey region) can be seen surrounded by the exocrine tissue (dark grey). An automated registration procedure (Haring et al., 2017) was done to overlay the fluorescence signal onto the EM images (Fig. 6C) such that the fluorescence signal is correlated at high resolution across the entire EM field of view (Fig. 6D & E). Additional details of how the correlative acquisition and reconstruction were done are provided in sections 6.4 & 6.5 respectively.

## Discussion

We have shown that the SNR of a 1 μs px^-1^ image subject to a bias potential outperforms that of a 20 μs px^-1^ unbiased image. This has important ramifications for large-scale and volume EM studies in which throughput is a primary concern. Due to practical limitations on time, it is often the case that large-scale EM studies are conducted on a single specimen. Negative potential bias facilitates comparison studies by allowing for multiple specimens to be acquired in the same timeframe that would otherwise be necessary for a single specimen. Experiments on specimens prone to electron beam irradiation damage are likewise facilitated as the same SNR can be achieved with a considerably smaller electron dose.

Our simulations show that BSE collection is enhanced by an effective increase of the detector numerical aperture—by applying the bias potential we increase the range of angular distributions of the BSEs able to be collected. However, this does not fully explain the extent of the increase in SNR observed experimentally. In particular, the simulations predict roughly a factor two increase in signal collection as the bias voltage is raised to our maximum of 3 kV, while our empirical measurements show SNR improvements of one to two orders of magnitude. This disparity can be explained in part by the electron gain factor of the detector. Sakic et al., 2011 (Sakic et al., 2011) shows that the signal generated in the detector by the incident electrons increases linearly with energy between 200 – 10000 eV. Thus, in addition to increasing the amount of collected BSEs, the bias potential also leads to signal enhancement via BSE acceleration. At low bias voltages the images appear to be dominated by one particular source of noise—which we suspect derives from the scanning electronics. Increasing the bias potential in this regime leads to an exponential rise in the SNR as this noise source is drowned out (Sup. Fig. 8). At sufficiently high bias voltages, the image noise is instead dominated by shot noise, stemming the exponential rise in SNR beyond 1.5 kV. The practical limit to the amount of bias potential we are able to apply is limited by the dielectric breakdown in vacuum. We estimate for our particular setup that the breakdown voltage occurs above 3 kV—well beyond the point at which the SNR plateaus.

Negative bias potential can be combined with imaging strategies currently in use for large-scale EM, such as the multi-scale approach taken by Hildebrand et al. (Hildebrand et al., 2017). The combination with an integrated microscope as demonstrated here could then offer a further benefit by in-situ selection of the regions of interest for high magnification acquisition. We envision a strategy in which regions of interest are first identified via fluorescence microscopy, then automatically navigated to and imaged with high resolution EM (Koning et al., 2019). Higher throughput could then be realized through a combination of faster acquisition via the negative bias potential, the removal of additional rounds of imaging, and the elimination of overhead from the entire imaging pipeline.

Further throughput enhancement could be obtained in several ways. One option would be to increase the current, thus increasing the per-pixel electron dose. However, doing so can come at the expense of resolution. Higher beam currents require larger aperture sizes which result in greater chromatic and spherical aberration. This can be problematic for many biological applications in which keeping aberrations at a minimum is critical for reaching a desired resolution, e.g. resolving neuronal connections, nuclear pores, or cell-cell junctions. Alternatively, the signal may be strengthened by increasing the landing energy. This may also be disadvantageous as too great a landing energy will result in partial transmission of electrons through the tissue section. In addition to reducing the number of generated BSEs in the tissue, this will increase the noise level by detection of accelerated BSEs generated in the underlying substrate. Finally, more signal could be generated by increasing the amount of staining material in the sample. This is a common approach for certain applications within large-scale EM such as neuronal connectomics, where an almost binary level of contrast may still be acceptable (Kuipers et al., 2015). Our stage bias approach holds promise to decrease acquisition times also in these applications, provided the lower limit imposed on dwell time by the detector response time is not reached.

## Material & Methods

### Modeling

All simulations were performed in Electron Optical Design (EOD) (Lencová and Zlámal, 2008). Descriptions of how the simulations were carried out are provided in the main text.

The angular distribution of signal electrons generated by a beam of primary electrons at a normal incident angle can be approximated by Lambert’s cosine law (Reimer, 1998). The probability of sampling a ray with angle *θ* to the normal of the surface is then proportional to cos *θ* sin *θ* = sin 2*θ*. If **U** is a random uniform distribution between 0 and 1, then

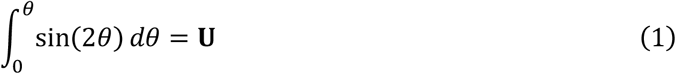

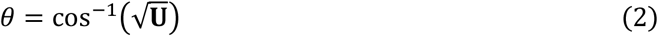

from which the initial angle of a signal electron can be chosen at random for use in simulations.

### Tissue and sample preparation

Fixed Rat pancreas tissue were post fixed for 2 hours in 1% osmiumtetroxide and 1.5% potassium ferrocyanide in 0.1 M cacodylate for 2hrs at 4°C. Followed by dehydration in a graded series of ethanol and finally embedded in epon. Ultrathin section of 80nm were cut and placed on ITO glass. Sections were blocked for 30 min with 1% bovine serum albumin (BSA; Sanquin, The Netherlands) in tris-buffered saline (TBS), pH 7.4. Next, anti-insulin (guinea pig; 1:50 in 1% BSA/TBS) was incubated for 2 h, followed by three washes of 5 min with TBS and subsequent incubation for 1 h with biotinylated secondary antibody (donkey-anti-guinea pig; 1:400 in 1% BSA/TBS, Jackson Immunoresearch, UK) followed by three washes in TBS. Finally, streptavidin conjugated Alexa594 (1:200, in 1% BSA/TBS, Life Technologies) was added for 1 h followed by three washes in TBS and two with MilliQ water. Hoechst staining was performed for 10min followed by a washing step with MilliQ water.

### Signal-to-noise measurements

The signal-to-noise ratio is calculated from computing the cross-correlation coefficient, *R_n_*, between successive scan lines, *I_i_*, and *I*_*i*+1_, of individual EM images following the method presented in (Joy, 2002). The cross-correlation coefficient is given by

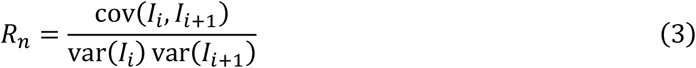

The signal-to-noise ratio is then calculated from

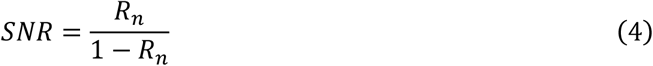

Additional SNR measurements were made based on the image power spectral density (PSD). The PSD is calculated by taking the squared magnitude of the 2D FFT. Assuming that the noise is white (uncorrelated from pixel to pixel), the PSD is equal to a constant (noise) term plus a varying (signal) term that dominates at low spatial frequencies. The noise term is measured by choosing a cutoff frequency above which the PSD is flat, and taking the magnitude of the PSD in this flat area. The noise variance is given by the integral of the PSD under the constant term. The signal variance is given by the total integral under the PSD minus the noise variance. The signal-to-noise ratio is then computed as the ratio of the variances.

### Integrated microscopy workflow

Fluorescence microscopy was done in the integrated microscope via the Delmic SECOM (Delmic B.V.), which has been retrofitted into the vacuum chamber of an FEI Verios SEM (Thermo Fisher Scientific) such that the two microscopes share a common sample stage and optical axis (Liv et al., 2013; Zonnevylle et al., 2013). With this configuration we are able to achieve sub 10 nm overlay precision without a reliance on fiducial markers or manual input (Haring et al., 2017). The SECOM was equipped with a CFI S Plan Fluor ELWD 60XC microscope objective (Nikon), which was chosen for its high magnification in combination with an extra-long working distance (2.60 - 1.80 mm). This lens enabled greater bias potentials to be reached without risking electrical breakdown in vacuum—at the cost of a somewhat lower numerical aperture (0.70 NA). Each FM image was comprised of two 3 s exposures: (1) 555 nm excitation for Alexa 594 labelling of insulin and (2) 405 nm excitation for the Hoechst counterstain.

An overview of imaging conditions is provided in Table 1. Fluorescence microscopy image tiles were acquired in a 4×3 grid encompassing the tissue section. Low magnification EM images of the same (but slightly smaller) field of view were acquired immediately following the acquisition of each FM image tile. An automated alignment procedure was then executed to register each set of FM and EM image pairs (Haring et al., 2017). The information necessary for registration was stored in the metadata of the image tiles for use in post-processing (6.5). The stage was then translated by 170 μm such that the FM images overlapped by a significant margin, whereas the low magnification EM tiles did not. This was done to prevent damage to the FM images due to e-beam irradiation. Following acquisition of the low magnification, correlative 4×3 grid, a 40×30 grid of high magnification EM image tiles was acquired over the section. Each image was acquired in immersion mode at 3 keV primary beam energy with a - 1.5 kV bias applied to the stage, resulting in a 1.5 keV landing energy. Of the 1200 high magnification EM images acquired, 113 were discarded as they consisted of only either epon or the substrate.

**Table 1:** Imaging parameters used for the full-section acquisition of 80 nm rat pancreas tissue via the integrated light-electron microscope.

|  | FM | EM (low mag) | EM (high mag) |
| --- | --- | --- | --- |
| Resolution | 107.8 nm px<sup>-1</sup> | 38.8 nm px<sup>-1</sup> | 4.86 nm px<sup>-1</sup> |
| Dwell | | 5 $\mu\text{s}$ | 5 $\mu\text{s}$ |
| Exposure | 3 s |  |  |
| Field of View | 220 $\mu\text{m}$ | 160 $\mu\text{m}$ | 20 $\mu\text{m}$ |

### Reconstruction

Following image acquisition, EM images were post-processed with histogram matching to correct for variations in intensity thought to have arisen from electron source drift during acquisition (variation in the bias potential delivered by the external power supply was negligible). No corrections were performed on the FM images. FM and EM image dataset was then uploaded to a local server running an instance of render-ws^1^. EM images were stitched together using the method presented in (Khairy et al., 2018). The correlative overlay between the FM and low magnification EM image tiles was done using the registration metadata collected at time of acquisition as described in 4.4.

The process of correlating the FM and stitched, high-magnification EM image tiles consisted of several steps. First, for each low-magnification EM tile, the set of overlapping high mag EM tiles was found. A composite image of the overlapping tiles was then rendered and processed with SIFT to find corresponding point matches with the low mag EM tile (Lowe, 1999). An affine transformation was then computed for this set of features and propagated to the FM tiles such that they overlaid precisely with the stitched together, high mag EM image tiles. The entire sequence of post-processing steps is compiled in a series of jupyter notebooks available in an online repository^2^.

Small 1024 × 1024 px^2^ images of the reconstructed dataset are then rendered and exported in a pyramidal format for visualization with CATMAID (Saalfeld et al., 2009). Within CATMAID, the FM images are given a false color transformation and the EM images are contrast-inverted for visualization purposes.

## CRediT authorship contribution statement

**Ryan Lane**: Conceptualization, Methodology, Investigation, Data curation, Writing – Original Draft. **Yoram Vos**: Conceptualization. **Anouk Wolters**: Resources. **Luc van Kessel**: Methodology. **Ben Giepmans**: Writing – Review & Editing. **Jacob Hoogenboom**: Writing – Review & Editing, Supervision, Project administration, Funding acquisition.

## Declaration of Competing Interest

R.L., Y.V., A.H.G.W., L.K., and B.N.N.G. declare that they have no competing interests. The integrated microscope used in this study is a product of Delmic BV. of which J.P.H. is a co-founder and shareholder.

## Acknowledgements

We thank Carel Heerkens and Ali Mohammadi-Gheidari for helpful discussions. This research is financially supported by the Dutch Research Council (NWO) through a Building Blocks of Life grant (737.016.010).

---

**Sup. Fig. 7.**
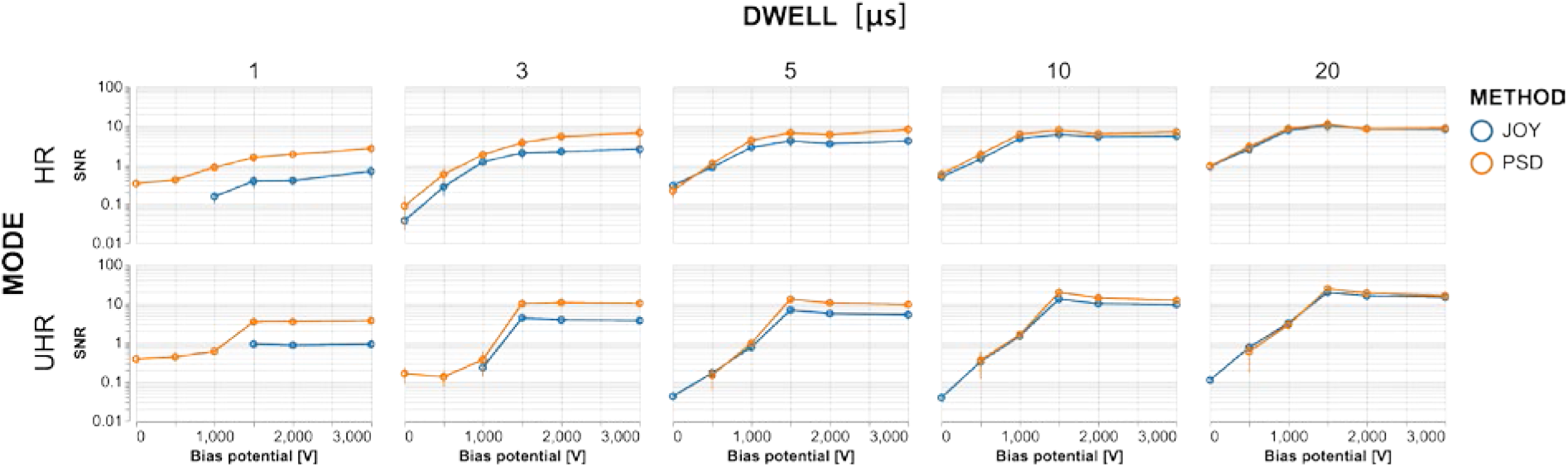
Independent SNR measurement methods show a high level of agreement, particularly for images with longer integration times. Measurements diverge somewhat for lower dwell images in which high frequency noise artefacts are more prominent. Both methods yield the more general trend of a steep increase up to 1.5 kV bias potential followed by a levelling off to 3 kV. Both methods also show a more drastic rise and plateau for UHR-mode, whereas the trend for HR-mode is more gradual.

**Sup. Fig. 8.**
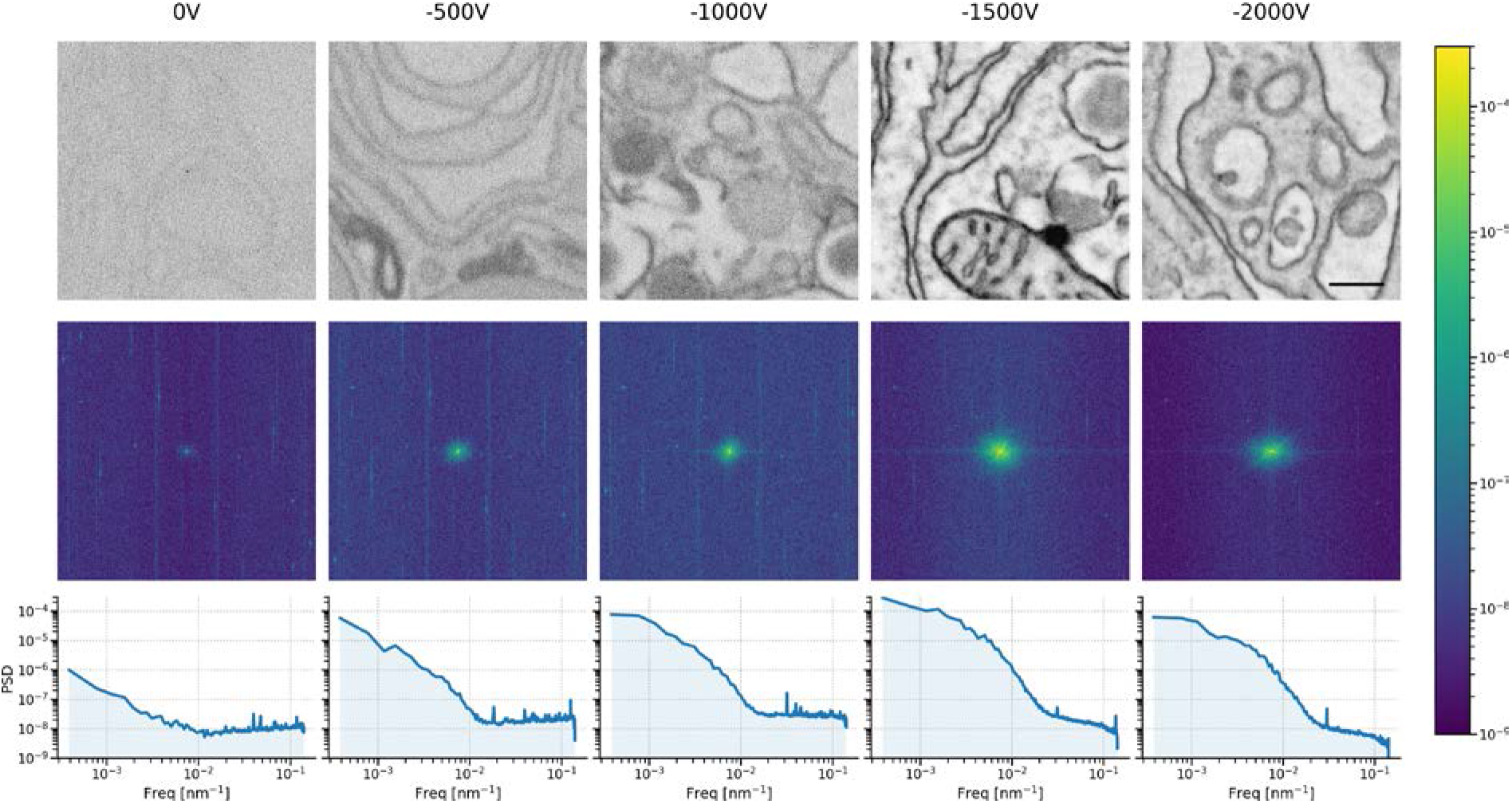
Noise contributions suspected to originate from the scanning electronics are suppressed with increasing bias potential. Top: sequence of 5 μs dwell tissue images acquired in immersion mode with varying amounts of stage bias. Center: 2D PSDs of tissue images showing the central spot, which represents most of the signal, becoming more prominent with increasing bias potential up to −1.5 kV. The 2D PSDs exhibit noticeable streak artefacts at higher frequencies, particularly in the lower bias potential images. We attribute these streaks to electric interference from the scanning electronics. Furthermore, there is a constant offset, which is likely a combination of shot noise from various sources, and may also include a component from the scanning electronics. Bottom: Azimuthally averaged power spectral density estimates show a division between the low frequency (primarily signal) and high frequency (primarily noise) portions of the tissue images. As the suspected scanning electronics noise is drowned out, the SNR improves dramatically. Scale bar: 500 nm.

## Footnotes

1 https://github.com/saalfeldlab/render

2 https://github.com/lanery/iCAT-workflow

## Notes

https://github.com/lanery/Lane-optimization-2020

http://www.nanotomy.org/OA/Lane2020/Lane.html

